# Small RNAs from the plant pathogenic fungus *Sclerotinia sclerotiorum* highlight candidate host target genes associated with quantitative disease resistance

**DOI:** 10.1101/354076

**Authors:** Mark C Derbyshire, Malick Mbengue, Marielle Barascud, Olivier Navaud, Sylvain Raffaele

**Affiliations:** Centre for Crop and Disease Management, Curtin University, Perth, WA, 6102, Australia; Laboratoire des Interactions Plantes Micro-organismes, INRA, CNRS, Université de Toulouse, Castanet Tolosan, France

## Abstract

Plant pathogenic fungi secrete effector proteins and secondary metabolites to cause disease. Additionally, some produce small RNAs (sRNAs) that silence transcripts of host immunity genes through RNA interference. The fungus *Sclerotinia sclerotiorum* infects over 600 plant species, but little is known about its molecular interactions with its hosts. In particular, the role of sRNAs in *S. sclerotiorum* pathogenicity has not been determined. By sequencing sRNAs *in vitro* and during infection of two host species, we found that *S. sclerotiorum* produces at least 374 highly abundant sRNAs. These sRNAs mostly originated from polymorphic repeat-rich genomic regions. Predicted gene targets of these sRNAs, from 10 different host species, were enriched for immunity-related functional domains. Predicted *A. thaliana* gene targets of *S. sclerotiorum* sRNAs were significantly more down-regulated during infection than other genes. *A. thaliana* gene targets were also more likely to contain single nucleotide polymorphisms (SNPs) associated with quantitative disease resistance. In conclusion, sRNAs produced by *S. sclerotiorum* are likely capable of silencing immunity components in multiple hosts. Prediction of fungal sRNA targets in host plant genomes can be combined with other global approaches, such as genome wide association studies and transcriptomics, to assist identification of plant genes involved in disease resistance.

## INTRODUCTION

Fungal phytopathogens largely rely on small secreted proteins, termed effectors, to infect and cause disease (1). Effectors can enter plant cells or act in the apoplast to manipulate host cell functions and promote fungal invasive growth (2).Necrotrophic fungi, which actively kill host cells, may also secrete various secondary metabolites to facilitate plant colonization (3). Recently, small RNAs produced by the plant pathogenic fungus *Botrytis cinerea* were also shown to be transferred into host tissues, where they silence host genes to facilitate infection (4). Subsequent studies have characterised a further sRNA from *B. cinerea* and sRNAs from the wheat rust fungus *Puccinia striiformis f. sp. tritici* with host-gene silencing properties (5, 6).

sRNAs are short non-coding sequences of RNA that are usually between 20 and 30 nucleotides long (7). Complementary base pairing of sRNAs with homologous mRNA sequences guides a group of proteins, called the RNA induced silencing complex (RISC), that mediate mRNA degradation or inhibition of translation (8). This process is known as RNA silencing, and it is involved in various cell functions such as development, transcription, translation and defence against viruses and transposable elements (7). In *B. cinerea*, a single sRNA potentially has the ability to target 15 genes in *Arabidopsis thaliana*, including WRKY transcription factors, receptor-like kinases and cell wall modifying enzymes (5). Many of the host genes targeted by the small RNAs of pathogenic fungi discovered to date exhibit functional domains typically associated with plant immune responses. In this way, fungal pathogen sRNA function may be analogous to that of pathogen effector proteins.

Another feature of sRNAs that makes them similar to effectors is their association with repetitive sequences. Many sRNAs are, in fact, often directly transcribed from transposable elements (7). The genomes of several filamentous pathogens have evolved towards compartmentalisation into repeat-rich, gene sparse regions that contain effector genes, and repeat-poor gene-rich regions that contain housekeeping genes (9, 10). Although there have been several sRNA profiling studies on plant pathogenic fungi (11–14), whether sRNA loci are associated with repeat-rich, gene poor regions of fungal genomes has not yet been considered.

Plant resistance to several fungal pathogens relies on the recognition of a single fungal effector by a plant resistance protein in a ‘gene-for-gene’ manner (15, 16). This leads to simple segregation between fully susceptible and fully resistant plants in host populations. However, infection by *S. sclerotiorum* results in a gradient of resistance phenotypes, controlled by a complex genetic program designated as quantitative disease resistance (QDR) (17).

Because QDR relies on numerous small-effect loci, unravelling the molecular bases of QDR is a major challenge in plant pathology. To identify portions of the genome containing markers linked with the QDR response, studies have used association genetics approaches (18, 19) such as genome wide association study (GWAS) (20). In the context of plant disease resistance, GWASs involve subjecting a diverse group of natural plant accessions to the same disease pressure. Linear models are then used to assess the predictive power of genomic markers for the level of disease whilst accounting for population structure (21–23). However, GWASs are limited by their ability to detect rare alleles and identify causative mutations associated with traits controlled by a large number of small effect loci (24). To circumvent these limitations, GWASs can be combined with other approaches such as biparental QTL mapping (25) and RNA sequencing (26). Since sRNAs from plant pathogenic fungi are likely to suppress host genes functioning in disease resistance, we hypothesised that the prediction of plant genes targeted by fungal sRNAs could be combined with a GWAS to aid identification of plant genes relevant to QDR.

Like *B. cinerea*, *Sclerotinia sclerotiorum* is a plant pathogen belonging to the *Sclerotiniaceae* family of Ascomycete fungi, which is able to infect hundreds of plant species (27, 28). *S. sclerotiorum* is widely dispersed throughout the world and poses a significant threat to agricultural production (29). A finished genome for *S. sclerotiorum* strain 1980 is available (30) and GWAS of QDR to *S. sclerotiorum* in *A. thaliana* has been reported (26). A previous study has shown that *S. sclerotiorum* produces sRNAs *in vitro* (11). Whether it also produces sRNAs during plant infection, and the potential roles of these sRNAs in plant infection are, thus far, undescribed.

We found that *S. sclerotiorum* produces 374 highly abundant sRNAs during infection of its two host species *Phaseolus vulgaris* and *A. thaliana*. In *A. thaliana*, predicted target genes were more likely to be significantly down-regulated during infection than other genes. Among the *A. thaliana* targets, there were significantly more genes associated with QDR by GWAS than expected by chance. Our data indicate that plastic regions of the *S. sclerotiorum* genome generate transposable element-derived sRNAs that potentially target numerous plant genes likely involved in QDR. They also suggest that fungal sRNA sequencing and identification of their plant targets can be complementary to other available approaches for the search of genes controlling plant QDR.

## MATERIAL AND METHODS

### Fungal cultures and inoculation of host plants

*Sclerotinia sclerotiorum* isolate 1980 was grown on potato dextrose agar plates for 4 days at 24°C. *Arabidopsis thaliana* accession Col-0 and *Phaseolus vulgaris* genotype G19833 were grown for 4 weeks before inoculation, under controlled conditions at 22°C, under a light intensity of 120 μmol/m2/s for 9 hours per day. Five mm-wide plugs containing actively growing *S. sclerotiorum* mycelium were inoculated onto fully developed leaves. Inoculated plants were placed in Percival AR-41 chambers at 80% humidity under the same day/light conditions as for plant growth for the duration of the infection. Samples for RNA extraction were harvested as described in (26), separating the centre and periphery of 2.5 mm wide disease lesions with a scalpel blade.

### RNA extraction and sequencing

Small RNAs were extracted and sequenced as described in (31). Briefly, samples harvested *in planta* and from *S. sclerotiorum* grown *in vitro* in Potato Dextrose Broth were processed using the NucleoSpin miRNA kit (Macherey-Nagel) following the instructions of the manufacturer. Small RNA sequencing was performed by Fasteris SA (Switzerland) on a HiSeq 2500 instrument using 50 bp single-reads.

### Quality filtering and mapping of sRNA reads

Reads were quality filtered using Trimmomatic version 0.22 (32) with the settings ‘ILLUMINACLIP:Adapters.fasta:2:1:1’, where ‘Adapters.fasta’ is a file containing adapter sequences, and their reverse complements, for the Illumina small RNA library prep kit. After quality filtering, reads from all samples were subjected to a further alignment-based filtering procedure. Bowtie version one with the settings ‘-v 0’ and ‘-a’, to obtain all exact genomic matches, was used for each step. For *A. thaliana*, reads from all samples that mapped to the sense strand of reads in the mock sample were discarded (antisense alignments were not discarded as small RNA sequencing is stranded). Then, reads that mapped to the sense strand of host gene transcripts, host gene non-coding RNAs downloaded from Rfam (version 13.0) and *S. sclerotiorum* non-coding RNAs downloaded from Rfam were discarded sequentially. Only reads that passed all of these steps and that were subsequently exactly mapped to the reference genome of *S. sclerotiorum* 1980 (30) were kept. The same procedure was followed for *P. vulgaris* using sRNA sequences predicted for this species obtained by (33). For differential expression analysis, the two filtering procedures for the two different hosts were both applied to the *in vitro* sample to create two different filtered *in vitro* datasets for analysis. This controlled for differences in filtering procedures that may cause artificial observation of changes in sRNA expression.

### Differential expression analyses of small RNAs

Differential expression analysis was performed using DESeq version 1.22.11 (34). Each unique sRNA was considered a single entity with a raw read count in this analysis. All *in planta* samples were compared with *in vitro* samples using a negative binomial test. Unique reads that exhibited a P adjusted value of below 0.05 were considered differentially expressed.

### Small RNA target prediction

To predict targets of *S. sclerotiorum* sRNAs, we used the psRNATarget online server (35). For the over-representation test of GO terms in *A. thaliana,* we used an E value of <= 3 resulting in 1,368 predicted sRNA target genes (1,789 including alternately spliced transcripts). For all other tests, we used a more stringent E value of <= 2.5 resulting in 408 predicted sRNA target genes (539 including alternately spliced transcripts). We used the same technique to predict targets of *B. cinerea* sRNAs in the *A. thaliana* genome. All other parameters were default. For all species in which we predicted sRNA targets, we first compared protein sequences against RepBase version 23.05 (36) using BLASTp. Any mRNAs whose proteins were homologous to sequences in RepBase with an e value of <= 1.0e^−10^ were not used in target prediction. This is because these proteins are likely host transposable element genes, which could be similar to fungal sRNAs by virtue of their shared evolutionary origins.

### Test for over-representation of functional domains

To test for over-representation of Gene Ontology (GO) terms among *A. thaliana* predicted targets of *S. sclerotiorum* sRNAs, we used the R bioconductor package TopGO (37). We performed the test separately for the molecular function and biological process categories of GO terms. In each instance, we used terms with 5 or more annotated genes. P-values for GO and PFAM enrichment were obtained using Fisher’s exact test and adjusted with the Benjamini-Hochberg correction (38). We also predicted targets in 10 different host transcriptomes from Phytozome. These included all species highlighted in supplementary figure 1. In all of these species, we performed an over-representation test of PFAM domains. To do this, we first created a list of all PFAM domains within the predicted annotations. We then created a list of PFAM domains present in predicted targets of sRNAs. The proportion of each PFAM domain in each set, either sRNA targets or non-targets, was compared using Fisher’s exact test. The PFAM domains with a p value of < 0.05 and an increased proportion in sRNA target genes were considered significantly over-represented.

### Messenger RNA sequencing and differential expression analysis

The expression of *A. thaliana* Col-0 genes during *S. sclerotiorum* infection corresponds to expression at the edge of necrotic lesions relative to non-challenged plants as reported in (26), data downloaded from the NCBI Gene Expression Omnibus (accession GSE106811). To determine expression of *A. thaliana* genes during *B. cinerea* infection, we obtained a dataset from (39) from the NCBI sequence read archive (https://www.ncbi.nlm.nih.gov/sra). The accessions were SRX1705130, SRX1705129 and SRX1705128, mock inoculated plants and SRX1728, SRX1729, SRX1730, 24 hours post inoculation with *B. cinerea.* DESeq2 version 1.10.1 was used to assess log_2_(fold change) between the mock-inoculated and *B. cinerea* infected samples.

### Test for down-regulated expression of fungal sRNA target genes in *A. thaliana*

To test whether predicted targets of fungal sRNAs in *A. thaliana* were more likely to be down-regulated during infection, we assessed the log_2_(fold change) (LFC) in expression of these genes during fungal infection relative to controls. First, we compared the median LFC of all sRNA targets with non-targets using a Wilcoxon test. Second, we used a non-parametric randomisation test based on the tests described in (40). A randomisation in our test consisted of assigning the status of ‘sRNA target’ and ‘non-sRNA target’ to *A. thaliana* genes randomly and calculating the difference between mean LFC values (ΔLFC). For each randomisation, we calculated ΔLFC as the mean LFC of non-sRNA targets minus the mean LFC of sRNA targets. The p values obtained from the test were the number of times ΔLFC was lower than the actual mean ΔLFC between sRNA targets and non-targets. We performed this analysis on four data sets: 1) Predicted *A. thaliana* targets of *S. sclerotiorum* sRNAs identified in this study during infection with S. *sclerotiorum.* 2) Predicted *A. thaliana* targets of *B. cinerea* sRNAs identified in Weiberg et al., (2013) during infection with *S. sclerotiorum.* 3) Predicted *A. thaliana* targets of *S. sclerotiorum* sRNAs during infection with *B. cinerea.* 4) Predicted *A. thaliana* targets of *B. cinerea* sRNAs during infection with *B. cinerea*.

### Test for over-representation of significant quantitative disease resistance scores among sRNA targets

To determine whether targets of *S. sclerotiorum* sRNAs were more likely to be significantly associated with QDR, we used the previously assessed markers from (26). We considered only genes that contained SNP markers, and only the most significant (highest −log_10_(P)) SNP in each gene. First, we conducted a Wilcoxon rank sum test of difference in median −log_10_(P). Second, we used a similar randomisation test to the test used for down-regulated expression of sRNA targets. In this instance, each randomisation recorded the proportion of targets and non-targets that exhibited a −log_10_(P) value of above 1.3 (or an untransformed score of below 0.05). The score for each randomisation was the proportion of targets with a significant SNP minus the proportion of non-targets with a significant SNP. This was termed the Δ%HQS. The P value derived from this test was the proportion of randomisations with a Δ%HQS higher than the actual data. Third, we performed a Fisher’s exact test of over-representation to test whether genes with significant SNPs were over-represented among genes that were predicted *S. sclerotiorum* sRNA targets.

## RESULTS

### *Sclerotinia sclerotiorum* small RNAs exhibit a characteristic length distribution and 5’ uridine bias

To identify *S. sclerotiorum* sRNAs expressed during infection of host plants, we conducted sRNA sequencing on *S. sclerotiorum* growing *in vitro* and during infection of the two host species *A. thaliana* and *P. vulgaris*. All samples were collected in triplicate and both the centre and border regions of disease lesions were harvested *in planta.* After adapter trimming and quality filtering, we obtained a total of 112,835,544 *in planta* reads across 12 samples and 21,394,528 *in vitro* reads across three samples. We filtered out reads that matched exactly (i) the sense strands of plant transcripts (either *P. vulgaris* or *A. thaliana*), (ii) plant non-coding RNAs (either *P. vulgaris* or *A. thaliana*), (iii) plant sRNAs (either a mock inoculated *A. thaliana* sample or a *P. vulgaris* sRNA data set obtained from (33)), (iv) the sense strands of *S. sclerotiorum* transcripts, (v) *S. sclerotiorum* non-coding RNAs from Rfam. The *in vitro* samples were filtered twice, once using *A. thaliana* sequences for steps (i)-(iii) and once using *P. vulgaris* sequences for steps (i)-(iii) to ensure that samples had undergone the same filtering procedures for differential expression analysis. The filtering process resulted in 14,798,914 mapped reads from all *in planta* samples, 635,493 *in vitro* reads when filtered against *A. thaliana* and 756,907 *in vitro* reads when filtered against *P. vulgaris*.

To determine the origin of sRNA reads in our samples, we analysed size distribution and 5’ nucleotide bias (Figure 1). Non-specific RNA degradation would result in uniform sRNA size distribution and random 5’ nucleotides. In contrast, >70% of raw reads were between 20 and 25 nucleotides long in all our samples. All *in planta* samples exhibited a peak in abundance at 22 nucleotides, whereas the *in vitro* sample exhibited equal proportions at 22 and 23 nucleotides (Figure 1 A). We also observed a bias toward uridine as the 5’ nucleotide in *S. sclerotiorum* sRNA reads (Figure 1 B). This bias was most pronounced for the 22 and 23 nucleotide reads in all samples. These data show that the sRNAs mapping to the *S. sclerotiorum* genome exhibited characteristics commonly attributed to sRNA biogenesis in diverse species.

**Figure 1.**
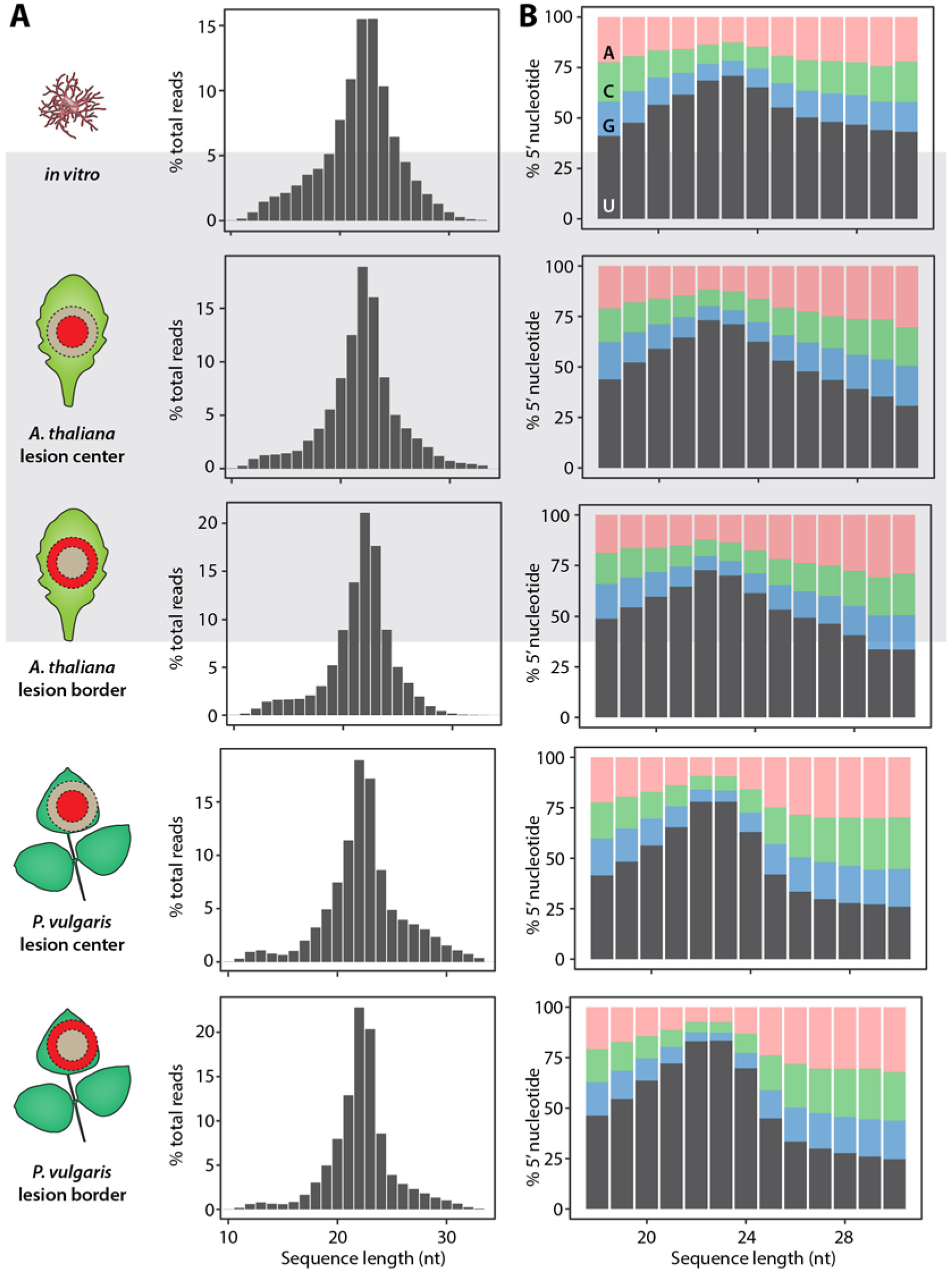
Length distribution and 5’ nucleotide bias of *S. sclerotiorum* sRNAs. (A) The percentage of reads (y axis) according to nucleotide (nt) sequence length (x axis) obtained *in vitro*, in *Arabidopsis thaliana* lesion centres and borders, and in *Phaseolus vulgaris* lesion centres and borders. (B) The percentage of <u>A</u>denine (pink), <u>C</u>ytosine (green), <u>G</u>uanine (blue) and <u>U</u>ridine (grey) in the 5’ prime position according to read length.

### Numerous *Sclerotinia sclerotiorum* sRNAs are highly expressed in two hosts

To identify *S. sclerotiorum* sRNAs that may be important for infection of host plants, we focused on sRNAs matching all of the following criteria: (i) corresponding to non-redundant sRNA sequences, (ii) of length of between 18 and 26 nucleotides, (iii) harbouring a 5’ uridine, and (iv) exhibiting over 100 reads per million *in planta*, in all three replicates collected from both host plants (either centre or border samples). This identified 374 abundantly expressed *S. sclerotiorum* core sRNA sequences in total (Figure 2A). Among these sRNAs, 322 were not significantly up-regulated during infection (Figure 2B), and 52 sRNAs (14%) were significantly up-regulated during plant infection relative to *in vitro* (Figure 2C). We did not find any sRNA reads that exhibited significant changes in abundance between the centres and borders of infection lesions (P adjusted > 0.05). To test whether the nature of the host plant affected the repertoire of sRNAs expressed by S. *sclerotiorum*, we considered sRNAs matching criteria i-iii above, and significantly up-regulated during infection of at least one host species relative to *in vitro* (Figure 2A). This identified a total of 94 sRNAs induced *in planta* (P adjusted < 0.05) (Figure 2D). Among these, 27 were up-regulated on *A. thaliana* only, 55 were up-regulated on *P. vulgaris* only, and 12 were shared between both hosts.

**Figure 2.**
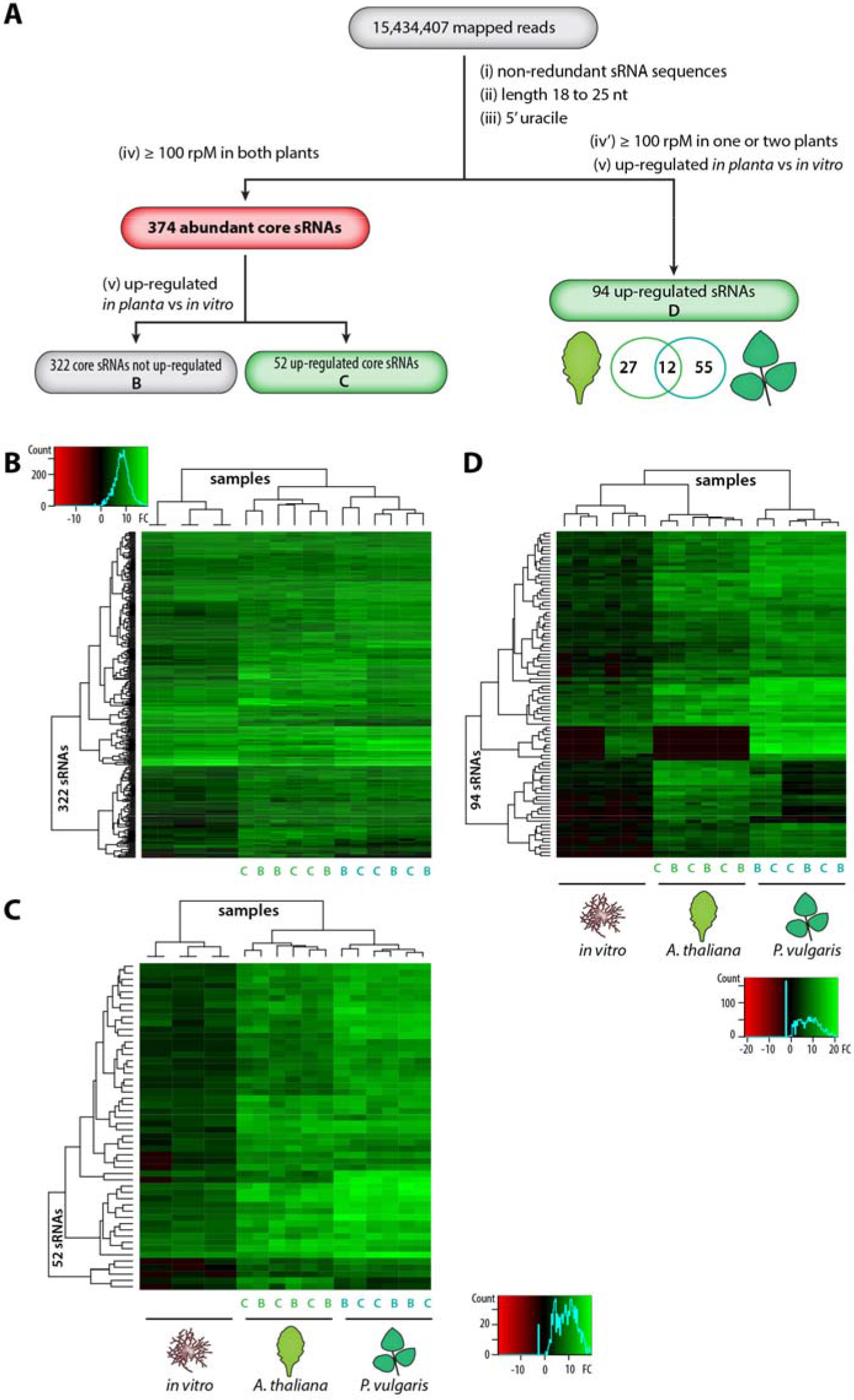
Identification of highly abundant fungal sRNAs and differential expression of fungal sRNAs *in planta*. (A) Five-step (i to v) pipeline used to identify the 374 highly abundant core sRNAs in *S. sclerotiorum* and differentially expressed sRNAs. Small RNAs expressed to a level of >= 100 reads per million (rpM) on both hosts in all replicates of at least one *in planta* sample (iv) were designated as abundant core sRNAs and were analysed for differential expression *in planta* relative to *in vitro* (v) (B, C). Small RNAs expressed over 100 rpM in at least one plant species (iv’) and up-regulated relative to *in vitro* (v) were analysed separately (D). (B-D) Heat maps of normalised expression data for the sRNAs identified using the procedure in (A). B, lesion border; C, lesion centre; FC, fold change; nt, nucleotides.

Together, these data show that a number of sRNAs are highly abundant *in planta* and many are significantly up-regulated during infection. Fungal cells residing at the centres and borders of disease lesions expressed similar sRNAs. In many cases, significant up-regulation of sRNAs occurred specifically in one host. This conclusion should be taken with caution considering the lack of methods explicitly designed to quantify sRNA expression and the relatively low coverage in our *in vitro* samples which may bias differential expression analysis. Therefore, we focused the following analyses on the 374 sRNAs highly abundant during the infection of both *A. thaliana* and *P. vulgaris*.

### *Sclerotinia sclerotiorum* sRNAs map to transposable element sequences and are in gene-poor polymorphic genome compartments

To determine loci from which the 374 highly abundant sRNAs originated in the *S. sclerotiorum* genome, we analysed overlapping annotations for genomic regions they mapped to, with a particular interest in the previously published REPET analysis of transposable elements in *S. sclerotiorum* (30). We found that these 374 sRNAs mapped to 3,676 loci, with 356 mapping to more than one locus and 18 mapping to a single locus. We found that 3,669 (99 %) of sRNA loci (including multiple mappings) were within 526 genomic regions annotated as transposable element sequence. The highest percentage of sRNA loci overlaps (32.5 %) were with long interspersed nuclear elements (LINEs, Figure 3 A). Only 7 (0.2 %) of the sRNA loci did not overlap with repeat sequences.

**Figure 3.**
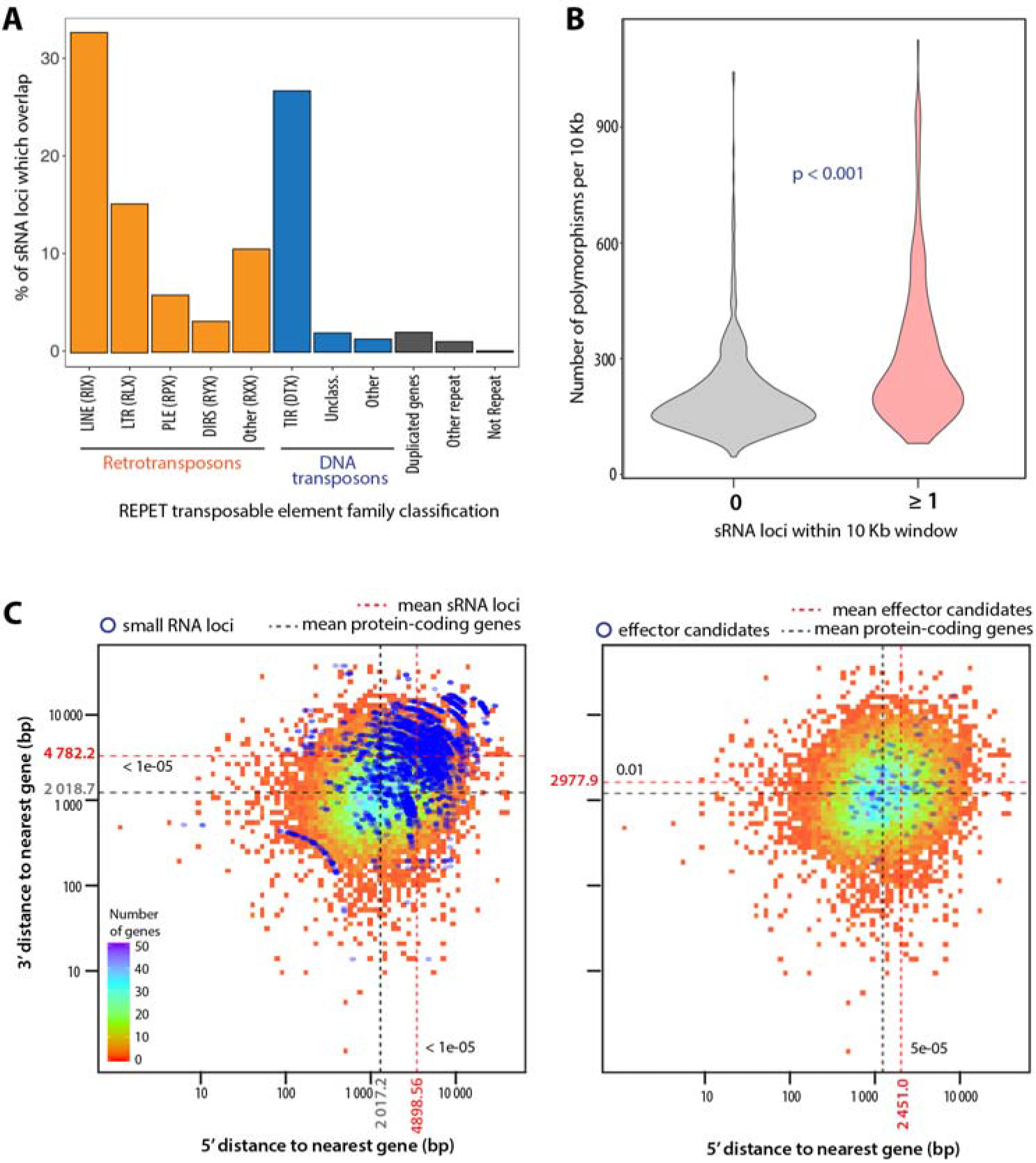
sRNA loci are associated with transposable elements, plastic and gene-sparse genomic regions. (A) Percentage of sRNA loci (y axis) that overlap different classes of repeat sequence annotated by REPET (x axis). Unclass., unclassified. (B) Number of polymorphisms (y axis) in 25 *S. sclerotiorum* isolates per 10 Kb sliding window. Ten Kb windows were analysed across the whole *S. sclerotiorum* genome and split into windows containing 0 and >= 1 sRNA locus (x axis). P-value for a Wilcoxon’s rank sum test is shown. (C) Distance to neighbouring genes in base pairs (bp) from the 5’ (x axis) and 3’ (y axis) ends of sRNA loci (left) and effectors (right). Blue points represent sRNAs or effectors and the underlying heatmap is for all *S. sclerotiorum* gene annotations. Mean distances for all *S. sclerotiorum* genes are show by grey dashed lines, mean distances for sRNAs or effectors are shown by red dashed lines represent with p-value of a Wilcoxon’s test for significant difference.

To determine whether the sRNA loci were more polymorphic than other regions of the genome, we assessed the number of polymorphisms in 10 Kb sliding windows throughout the genome from a panel of 25 *S. sclerotiorum* isolates (BioRxiv: doi/xxx). We counted polymorphisms if at least one individual did not exhibit the reference genome allele. The median number of polymorphisms was 235 for windows including at least one sRNA locus, but only 174 for windows not containing sRNA loci (P < 0.001, W = 54084, Wilcoxon’s rank sum test) (Figure 3 B). Strikingly, the proportion of windows showing 300 polymorphisms or more was 36% for windows including sRNA loci but only 12% otherwise. Because accurate mapping of short sequence reads to repetitive regions is challenging, we used variants that were filtered based on excess coverage and quality (BioRxiv: doi/xxx). Nevertheless, the excess of polymorphisms at sRNA loci, although robust, could have been over-estimated.

To determine the extent to which sRNA loci reside in gene-sparse regions, we calculated the distance between the 5’ and 3’ ends of all sRNA loci and the nearest gene borders, and for all *S. sclerotiorum* protein-coding genes, including predicted effector genes from (30) (Figure 3C). The mean 5’ distance to nearest gene was 4898.56 bp for sRNA loci and 2017.2 bp for protein-coding genes (P < 2.2e-16, W = 3053600). Similarly, the mean 3’ distance to nearest gene was 4551.631 bp for sRNA loci and 2018.7 bp for protein-coding genes (P < 2.2e-16, W = 31125000). The genes encoding effector candidates also resided further from the nearest gene than other protein-coding genes, albeit to a lesser extent than sRNA loci. The mean distance to the nearest gene was 2451.0 bp for effector candidate genes on the 5’ side (P = 0.01005, W = 442360) and 2977.9 bp on the 3’ side (P = 5.37e-05, W = 480430). Together, these data indicate that *S. sclerotiorum* sRNAs are derived from gene-sparse repetitive regions that are more polymorphic than the rest of the genome.

### Predicted plant targets of *Sclerotinia sclerotiorum* sRNAs are enriched for immunity-related functional domains

Several Ascomycete fungi produce sRNAs capable of modulating gene expression in their host plants (41). To identify predicted targets of the 374 highly abundant *S. sclerotiorum* sRNAs in plant genomes, we used psRNATarget (35). We performed both GO term and PFAM domain enrichment analyses using Fisher’s exact test. We predicted in the *A. thaliana* genome 1,368 sRNA target genes with an (E) value ≤ 3, among which 408 had an (E) value ≤ 2.5. Among gene functions strongly enriched in sRNA targets (Figure 4, Supplementary Tables 1 and 2), several were related to signalling (e.g. GO:0007165, GO:0019199, PF00069 and PF03107) and specifically plant immunity signalling (GO:0019199 and PF13855). Annotations related to plant defense responses (GO:0006952, GO:0052542, and PF12662), were also enriched. Other annotations enriched among predicted *S. sclerotiorum* sRNA targets included ontologies and domains related to hormone metabolism (GO:0009737, GO:0046345, GO:0008742) and redox metabolism (GO:0055114, GO:0019825, PF00175).

**Figure 4.**
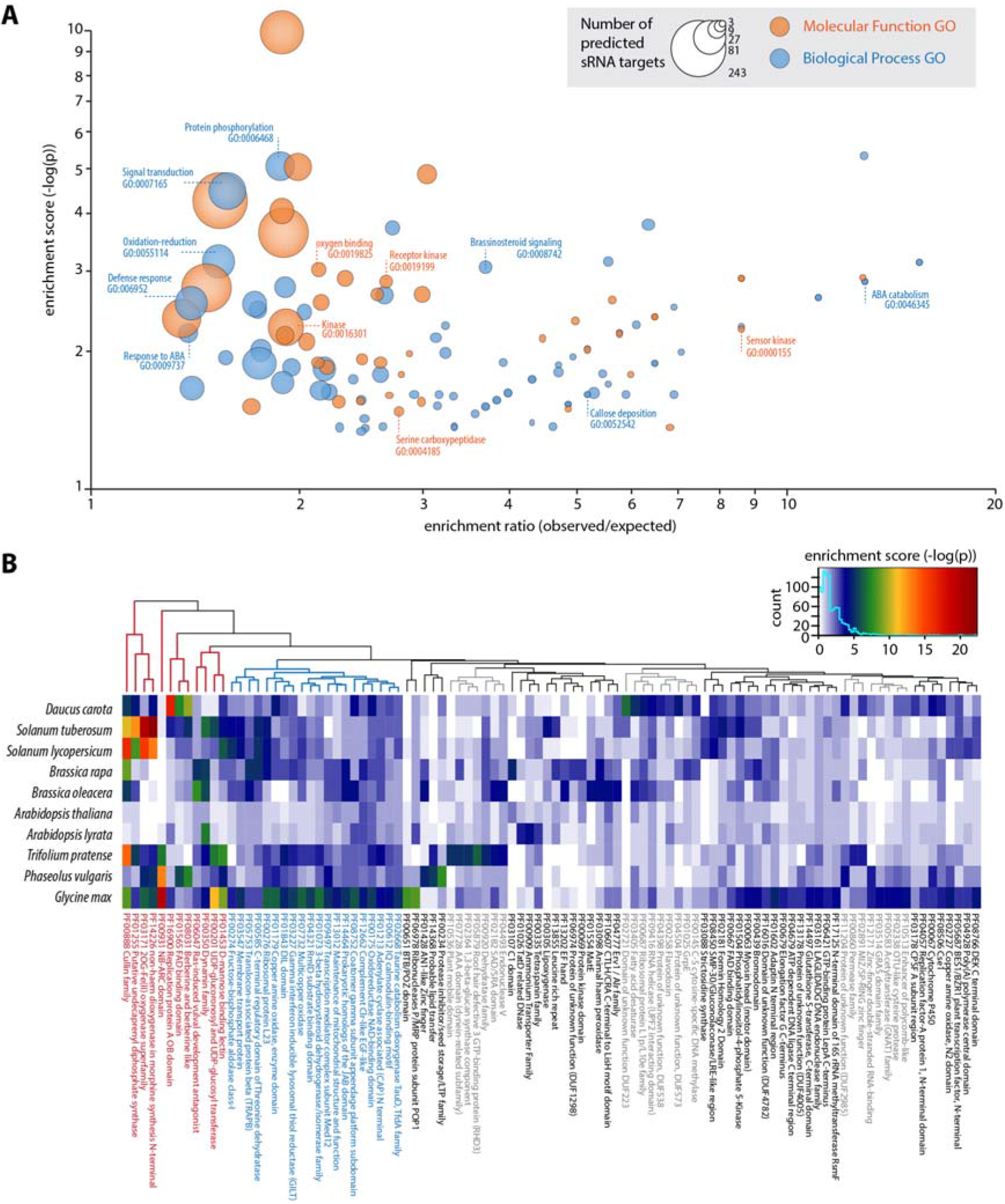
Enrichment of functional domains in plant targets of *Sclerotinia sclerotiorum* sRNAs. (A) Enrichment (−log(p) (y axis)) of Gene Ontology (GO) terms in putative *A. thaliana* targets of the 374 *Sclerotinia sclerotiorum* core sRNAs. The ratio of the observed to expected proportions of the GO terms in sRNA targets is on the x axis. The size of points represents the number of predicted sRNA targets annotated with the GO term. Molecular Function GO terms are in orange and Biological Process GO terms are in blue. Several terms discussed in the text are highlighted that indicate roles in signalling, hormone metabolism and defence against pathogens. (B) PFAM domains significantly enriched among putative gene targets of the 374 *S. sclerotiorum* core sRNAs in at least one of 10 plant species. The enrichment score is plotted as a heatmap, from white (−log(p) of 0) to dark red (−log(p) > 20). PFAMs enriched in most plants are labelled in blue, PFAMs strongly enriched in a few plants only are labelled in red.

*S. sclerotiorum* is able to infect hundreds of plant species, including carrot (*Daucus carota*), tomato (*Solanum lycopersicum*), rapeseed (*Brassica rapa*) and soybean (*Glycine max*). To determine the range of potential targets of *S. sclerotiorum* sRNAs in diverse host species, we predicted the targets of the 374 abundant *S. sclerotiorum* sRNAs in the genomes of 10 plant species using psRNATarget and analysed PFAM domains enriched for sRNA target genes. We found that all of these genomes exhibited numerous potential sRNA targets, on average 521 per host genome (Supplementary Table 3). A total of 98 non-redundant PFAM domains were enriched in *S. sclerotiorum* sRNA target genes in at least one plant species (Fisher’s exact test P < 1e^−04^) (Figure 4B). A number of these PFAM domains also indicated possible roles in plant immunity, such as PF13855, PF00931, PF01453, PF00305. Twenty PFAM domains were enriched among *S. sclerotiorum* sRNA targets in most plant species (Figure 4B, blue), notably including domains related to redox functions, secretion and transport and transcription. Despite being identified using the same set of 374 *S. sclerotiorum* sRNAs, different functional terms were over-represented in different host species. In particular, twelve PFAM domains were strongly enriched among *S. sclerotiorum* sRNA targets in a few species only (Figure 4B, red), such as PF00931 in legumes, PF00350 in *Brassicaceae* and potato, and domains related to the biosynthesis of secondary metabolites (PF01255 and PF14226) in *Solanaceae*. Overall, these data suggest that *S. sclerotiorum* sRNAs are able to target numerous host genes in phylogenetically diverse species. Many of the predicted domains in these target genes have been previously associated with disease resistance in plants.

### Predicted targets of *Sclerotinia sclerotiorum* sRNAs in *Arabidopsis thaliana* are more likely to be down-regulated during infection

To support host gene silencing by *S. sclerotiorum* sRNAs, we compared the expression of *A. thaliana* genes putatively targeted by fungal sRNAs with that of non-target genes during fungal infection. We analysed predicted targets of *B. cinerea* sRNAs (4), predicted targets of *S. sclerotiorum* sRNAs and all other *A. thaliana* genes during infection with *B. cinerea* (39) (Figure 5A) or during infection with *S. sclerotiorum* (Figure 5B). We considered gene expression log_2_(fold change) (LFC) relative to uninoculated controls. During *B. cinerea* infection, *A. thaliana* targets of *B. cinerea* sRNAs predicted by (4) exhibited a lower median LFC (-0.064) than targets of *S. sclerotiorum* sRNAs (median LFC= -0.019) and other genes (median LFC= 0.013; Wilcoxon’s rank sum test P = 0.0989, W = 1862900). Compared to non-target genes, the difference in median LFC (ΔLFC) was -0.077 for *B. cinerea* sRNAs targets and -0.032 for *S. sclerotiorum* sRNA targets. Conversely, during *S. sclerotiorum* infection, predicted *A. thaliana* targets of *S. sclerotiorum* sRNAs identified in this study exhibited a significantly lower LFC (-0.84) than targets of *B. cinerea* sRNA (median LFC= -0.14) and other genes (median LFC= -0.41; P = 0.03358, W = 4238200). Compared to non-target genes, *S. sclerotiorum* sRNAs targets had a ΔLFC of -0.43 and *B. cinerea* sRNAs targets a ΔLFC of 0.27.

**Figure 5.**
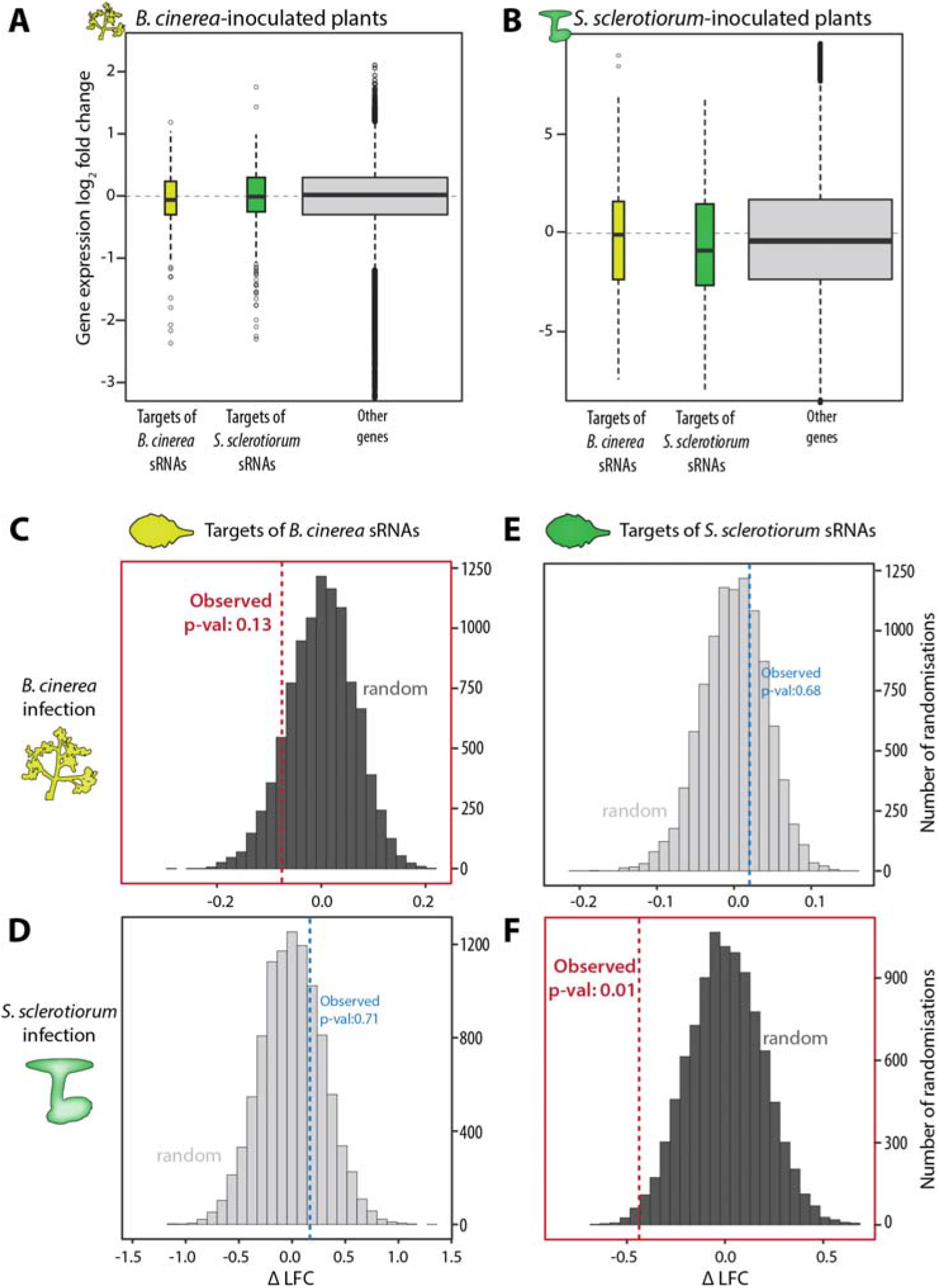
Predicted targets of *S. sclerotiorum* sRNAs in *A. thaliana* are significantly down-regulated during infection. (A) Expression Log_2_(fold change) (LFC) (y axis) upon inoculation by *Botrytis cinerea* for *Arabidopsis thaliana* genes that were predicted targets of *B. cinerea* sRNAs (yellow), predicted targets of *Sclerotinia sclerotiorum* sRNAs (green), and all other genes (grey). Horizontal black lines represent median LFC, whiskers represent interquartile range and boxes represent 2^nd^ and 3^rd^ quartiles. (B) The same as for (A) upon inoculation by *S. sclerotiorum*. (C-F) Distribution of difference in median LFC (ΔLFC) between genes targeted or not by sRNAs in 10,000 randomisations (grey). Dashed vertical lines represent the observed ΔLFC. Predicted *A. thaliana* targets of *B. cinerea* sRNAs during infection with *B. cinerea* (C) and infection with *S. sclerotiorum* (D). Predicted *A. thaliana* targets of the 374 abundant *S. sclerotiorum* sRNAs during infection with *B. cinerea* (E) and infection with *S. sclerotiorum* (F).

As a complementary analysis, we performed a randomisation to test for the likelihood of finding a lower ΔLFC in random *A. thaliana* gene sets than observed for fungal sRNA targets. During *B. cinerea* infection, only 13 % (equivalent to a one tailed P value of 0.13) of random *A. thaliana* gene samples had a lower ΔLFC than was observed from targets of *B. cinerea* sRNAs (Figure 5C), but 71% had a lower ΔLFC than targets of *S. sclerotiorum* sRNAs (~p=0.71, Figure 5D). Conversely, during *S. sclerotiorum* infection, 68% of random *A. thaliana* gene samples had a lower ΔLFC than targets of *B. cinerea* sRNAs (~p=0.68, Figure 5E), whereas only 1% had a lower ΔLFC than targets of *S. sclerotiorum* sRNAs (~p=0.01, Figure 5F). Together, these analyses suggest that *S. sclerotiorum* sRNAs could have a negative impact on the expression of their *A. thaliana* target genes during infection.

### Predicted targets of *Sclerotinia sclerotiorum* sRNAs in *Arabidopsis thaliana* are more likely to be associated with quantitative disease resistance

To support a role in disease resistance for predicted targets of *S. sclerotiorum* sRNAs in *A. thaliana*, we tested for the association of these genes with a QDR phenotype to *S. sclerotiorum* measured in *A. thaliana* natural accessions. For this, we exploited the probability of association with QDR against *S. sclerotiorum* for 204,648 single nucleotide polymorphisms (SNPs), obtained through a genome wide association study (GWAS) in 84 European accessions of *A. thaliana* (26). We identified 122,034 SNPs distributed in 23,688 gene models, and determined a score of association per gene (−log_10_ of the p-value for the most significant SNP for each gene). First, we found that the median score of association for predicted targets of *S. sclerotiorum* sRNAs was 0.82, significantly higher than the median score for non-targets (median = 0.67, Wilcoxon’s rank sum test P = 4.872e^−05^, W = 4159300) (Figure 6A). Genes with an association score >1.3 (corresponding to a pvalue < 0.05) represented 18.7% of *S. sclerotiorum* sRNA targets, but only 14.9% of other genes (Difference in proportion of high QDR score Δ%HQS= 3.8), Fisher’s exact test P=0.04). Second, we performed a randomisation test to determine the likelihood of having a Δ%HQS ≥ 3.8 in random *A. thaliana* gene sets of the same size (Figure 6B). We obtained a Δ%HQS < 3.8 in 98.1 % of randomisations indicating that *S. sclerotiorum* sRNA targets are significantly enriched in genes with an association score >1.3 (P = 0.0191).

**Figure 6.**
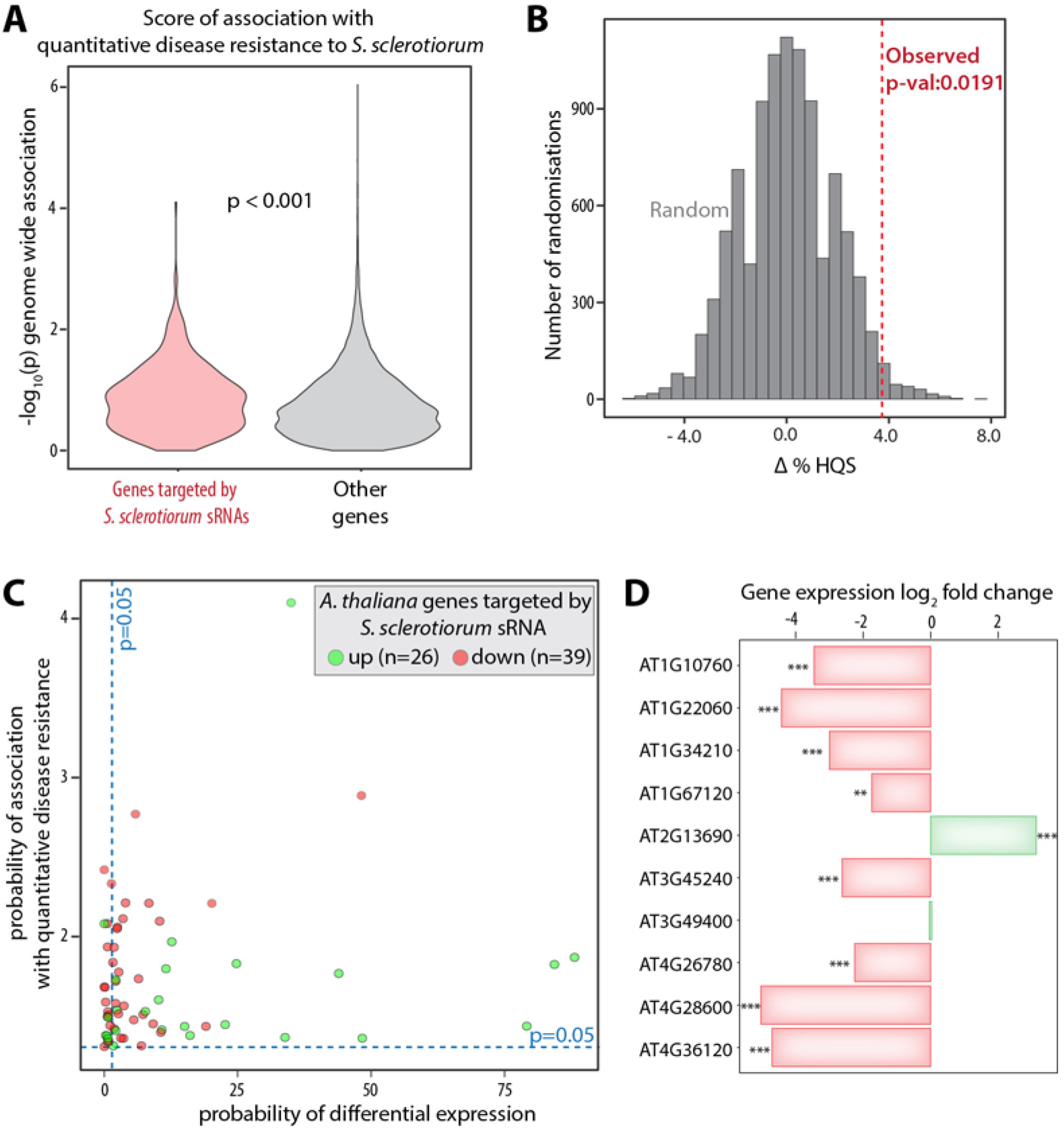
Association between high quantitative disease resistance (QDR) score and decreased expression during infection for *A. thaliana* genes targeted by *S. sclerotiorum* sRNAs. (A) Quantitative disease resistance (QDR) scores (−log10(P) of association) (y axis) for *A. thaliana* genes predicted to be targeted by the 374 abundant *Sclerotinia sclerotiorum* sRNAs (left) and other genes (right). P-value for a Wilcoxon’s test is shown. (B) Distribution of Δ%HQS (difference in the % of genes with a QDR association score>1.3 between genes targeted or not by sRNAs) in 10,000 randomisations (grey). The vertical dashed line represents the observed Δ%HQS for *A. thaliana* genes targeted or not by *S. sclerotiorum* sRNAs. (C) Distribution of *A. thaliana* genes with a QDR score above 1.3 (P < 0.05) that were predicted targets of *S. sclerotiorum* sRNAs, according to their QDR score (y-axis) and p-value for differential expression upon S. sclerotiorum inoculation (x axis). Genes up-regulated during infection are shown in green, genes down-regulated are shown in red. (D) Expression Log_2_(fold change) for *A. thaliana* genes targeted by *S. sclerotiorum* sRNA with the top 10 QDR scores. Eight genes were significantly down-regulated (red bars, *** = P < 0.001), one was significantly up-regulated (green bar, *** = P < 0.001).

To identify the most relevant targets of *S. sclerotiorum* sRNAs for plant resistance, we focused on the 65 *A. thaliana* predicted target genes showing (i) p<0.05 for differential expression upon *S. sclerotiorum* inoculation and (ii) p<0.05 for association with quantitative disease resistance in GWAS (Figure 6C). In agreement with our previous analysis (Figure 5), the majority of the 65 genes (39 genes, corresponding to 60%) were down regulated during *S. sclerotiorum* infection. Among the top 10 scores for association with QDR, 9 genes belonged to the list of 65, including 8 genes significantly down-regulated (from −1.74 to −5.03 fold) one gene significantly up-regulated (*AT2G13690* encoding a PRLI-interacting factor of unknown function) and one gene that did not exhibit any change in expression (*AT3G49400* encoding a Transducin/WD40 repeat-like protein of unknown function) (Figure 6D, Table 1). Down-regulated genes included a MIDASIN1 homolog (*AT1G67120*) associated with the assembly of the 60S ribosomal subunit, a mitochondrial nucleotide exchange factor GrpE (*AT4G26780)* associated with tolerance to heat stress in *A. thaliana*, three genes encoding proteins of unknown function (*AT4G36120*, *AT4G28600*, *AT1G22060*) and three kinases. *AT3G45240* encodes a geminivirus Rep interacting kinase (GRIK1) activating SnRK1 (SNF1-related kinases), *AT1G34210* encodes the plasma membrane LRR receptor-like serine threonine kinase SOMATIC EMBRYOGENESIS RECEPTOR-LIKE KINASE 2 (SERK2) related to the PRR co-receptor BAK1, and *AT1G10760* encodes the α-glucan, water dikinase STARCH EXCESS 1 (SEX1) required for starch degradation. Mutants in *SEX1* are more susceptible to the fungal pathogen *Colletotrichum higginsianum* (Engelsdorf et al., 2016) supporting the view that *S. sclerotiorum* sRNA targets can serve as probes to identify plant genes relevant for quantitative disease resistance.

**Table 1.**
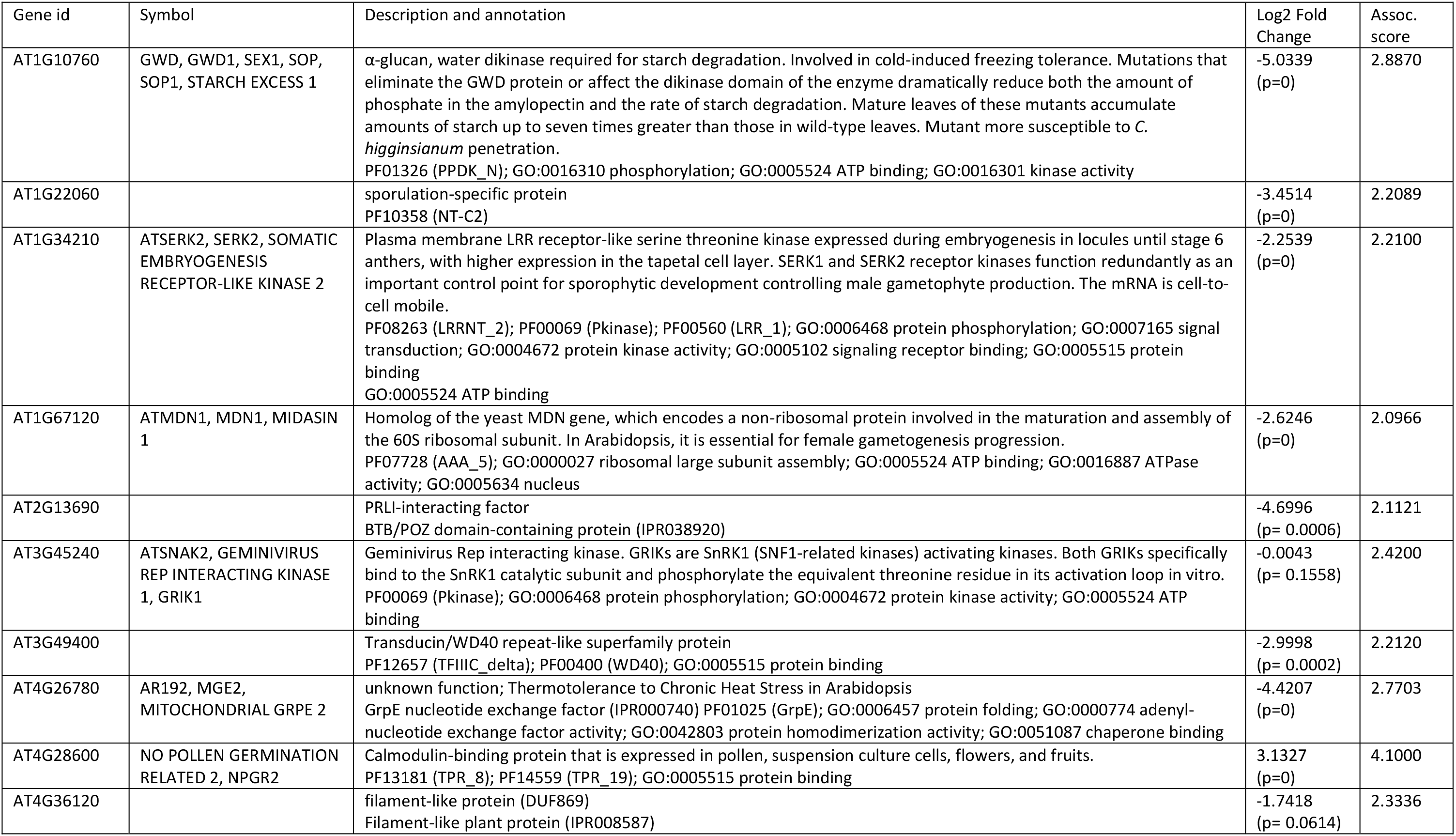
Predicted target genes of *S. sclerotiorum* sRNAs in *A. thaliana* with the top 10 scores for association with QDR.

## DISCUSSION

In this study, we demonstrate that the broad host range plant pathogen *S. sclerotiorum* exhibits numerous abundant sRNAs expressed on two different hosts. These sRNAs exhibit characteristics typical of previously described filamentous pathogen sRNAs, such as a length distribution peaking between 20 and 25 nucleotides and a 5’ uridine bias (13, 14, 42, 43). This echoes findings of (11) who analysed sRNAs produced by *S. sclerotiorum in vitro.* The 5’ uridine has been shown to be important for sRNA functions in distantly related species, such as *Drosophila melanogaster* and *A. thaliana* (44, 45). In these species, the presence of a 5’ uridine is important for directing the sRNAs to a specific Argonaut family protein, AGO1. This suggests that AGO1 (<u>sscle 03g027950</u>) is a key player mediating RNA silencing in S. *sclerotiorum.* Although plant AGOs have undergone extensive diversification compared with animal AGOs, most canonical plant miRNAs are incorporated into AGO1 clade proteins (46). Thus, the 5’ uridine of sRNAs may be an important feature conserved in evolution to facilitate trans-kingdom RNA silencing between plants and fungi. It is yet to be elucidated whether this bias in 5’ nucleotide exists among non-plant pathogenic fungi.

Most of these sRNAs mapped to repetitive genomic regions, suggesting that sRNAs, like some fungal effectors, may be strongly associated with transposable elements. The repeat-rich regions that contain effector genes often exhibit a reduced overall gene content. This phenomenon appears to be convergent among distantly related plant pathogens in both the fungal and oomycete classes (9, 47–50). In some species, effector genes within repeat-rich regions exhibit high levels of sequence diversification and tend to evolve more rapidly than the rest of the genome, an observation encapsulated in the ‘two-speed’ genome hypothesis (51). Clustering of repeats and effectors in specific genome niches probably results from selection against deleterious mutations in essential genes. Genome compartmentalisation seems most pronounced in fungi with a biotrophic or hemi-biotrophic component to their lifestyles, which specialise on one or a few host species (49, 50). In contrast, broad host-range necrotrophic fungal pathogens such as *S. sclerotiorum* and *B. cinerea*, exhibit relatively repeat poor genomes (30, 52). Nevertheless, effector genes may also be associated with repeat sequences in necrotrophic fungi and fungi with a broad host range (5, 53–55). Secretome analyses highlighted a number of candidate effector-like proteins in the genome of *S. sclerotiorum* (30, 56). Some of these predicted effector genes associate with repeat sequences. In this study, almost all sRNA loci were associated with transposable elements. Like effectors, *S. sclerotiorum* sRNA loci were, on average, further from gene sequences than genes were to each other. Although the number of polymorphisms in repetitive regions cannot be estimated with high accuracy, 10 Kb windows including sRNA loci were clearly more polymorphic than the rest of the genome. These findings indicate that *S. sclerotiorum* exhibits a fast-evolving, transposon-associated sRNA effector repertoire. The fact that sRNA loci were highly polymorphic would support the hypothesis that these regions are an important component of adaptive evolution to the very diverse host environments that populations of broad host range fungi encounter in nature. It is possible to envisage a scenario under which random targeting of sRNAs to host immunity genes in one or more species confers a selective advantage. A rapid turnover of sRNAs and large potential for inhibition of host genes through random sequence matches could thus create a high adaptive potential of *S. sclerotiorum* populations.

Many of the predicted host targets of the sRNAs identified in this study exhibited domains previously shown to function in plant immune responses (57, 58). Furthermore, upon infection with *S. sclerotiorum*, these target genes were far more likely to be down-regulated than non-targets. Although further functional assays would be required to draw firm conclusions, these data support the view that *S. sclerotiorum* actively suppresses host immunity genes with sRNAs.

To further understand the importance of trans-kingdom RNA silencing during plant infection with *S. sclerotiorum* we tested the degree of association with quantitative disease resistance for the predicted sRNA targets in *A. thaliana* exploiting a previous GWAS (26). The GWAS score was significantly higher in predicted *S. sclerotiorum* sRNA targets than non-targets. In the *A. thaliana* GWAS, the p-values of association with QDR remained, nevertheless, relatively modest for *S. sclerotiorum* sRNA targets (score ≤ 4.1, corresponding roughly to a false discovery rate of 1.0 e^−06^). Therefore, these genes may not have been selected as relevant for QDR based on GWAS data alone. However, the combination of GWAS, RNA sequencing, and fungal sRNA target predictions revealed novel candidate genes to be functionally characterised for association with QDR in the future. This combination of approaches may be useful in narrowing down candidate plant genes for functional characterisation in diverse plant-pathogen interactions, particularly in the absence of high resolution genetic maps and for species in which functional tests are challenging.

## ACCESSION NUMBERS

All RNA sequencing data have been deposited in the GenBank sequence read archive under BioProject number PRJNA477286

## SUPPLEMENTARY DATA

Supplementary Table 1: Tables1.xlsx

Supplementary Table 2: Tables2.xlsx

Supplementary Table 3: Tables3.docx

## FUNDING

This research was supported by funding from the Grains Research and Development Corporation and Curtin University [grant number CUR00023 to MCD], the European Research Council [grant number ERC-StG-336808 to SR] and the French Laboratory of Excellence TULIP [grant numbers ANR-10-LABX-41, ANR-11-IDEX-0002-02]. This study also benefited from support by the “visiting scientist” program of the TULIP Labex.

## CONFLICT OF INTEREST

The authors have no conflicts of interest to declare.

## REFERENCES

1 Lo Presti,L., Lanver,D., Schweizer,G., Tanaka,S., Liang,L., Tollot,M., Zuccaro,A., Reissmann,S. and Kahmann,R. (2015) Fungal Effectors and Plant Susceptibility. Annu. Rev. Plant Biol., 66, 513–545.

2 Toruño,T.Y., Stergiopoulos,I. and Coaker,G. (2016) Plant-Pathogen Effectors: Cellular Probes Interfering with Plant Defenses in Spatial and Temporal Manners. Annu. Rev. Phytopathol., 54, 419–441.

3 Friesen,T.L., Faris,J.D., Solomon,P.S. and Oliver,R.P. (2008) Host-specific toxins: effectors of necrotrophic pathogenicity. Cell. Microbiol., 10, 1421–1428.

4 Weiberg,A., Wang,M., Lin,F.-M., Zhao,H., Zhang,Z., Kaloshian,I., Huang,H.-D. and Jin,H. (2013) Fungal Small RNAs Suppress Plant Immunity by Hijacking Host RNA Interference Pathways. Science (80-.)., 342, 118–123.

5 Wang,Q., Jiang,C., Wang,C., Chen,C., Xu,J.-R. and Liu,H. (2017) Characterization of the Two-Speed Subgenomes of Fusarium graminearum Reveals the Fast-Speed Subgenome Specialized for Adaption and Infection. Front. Plant Sci., 8, 140.

6 Cingolani,P., Platts,A., Wang,L.L., Coon,M., Nguyen,T., Wang,L., Land,S.J., Lu,X. and Ruden,D.M. (2012) A program for annotating and predicting the effects of single nucleotide polymorphisms, SnpEff. Fly (Austin)., 6, 80–92.

7 Dang,Y., Yang,Q., Xue,Z. and Liu,Y. (2011) RNA Interference in Fungi: Pathways, Functions, and Applications. Eukaryot. Cell, 10, 1148–1155.

8 Pratt,A.J. and MacRae,I.J. (2009) The RNA-induced Silencing Complex: A Versatile Gene-silencing Machine. J. Biol. Chem., 284, 17897–17901.

9 Dong,S., Raffaele,S. and Kamoun,S. (2015) The two-speed genomes of filamentous pathogens: waltz with plants. Curr. Opin. Genet. Dev., 35, 57–65.

10 Fouché,S., Plissonneau,C. and Croll,D. (2018) The birth and death of effectors in rapidly evolving filamentous pathogen genomes. Curr. Opin. Microbiol., 46, 34–42.

11 Zhou,J., Fu,Y., Xie,J., Li,B., Jiang,D., Li,G. and Cheng,J. (2012) Identification of microRNA-like RNAs in a plant pathogenic fungus Sclerotinia sclerotiorum by high-throughput sequencing. Mol. Genet. Genomics, 287, 275–282.

12 Chen,R., Jiang,N., Jiang,Q., Sun,X., Wang,Y., Zhang,H. and Hu,Z. (2014) Exploring MicroRNA-Like Small RNAs in the Filamentous Fungus Fusarium oxysporum. PLoS One, 9, e104956.

13 Mueth,N.A., Ramachandran,S.R. and Hulbert,S.H. (2015) Small RNAs from the wheat stripe rust fungus (Puccinia striiformis f.sp. tritici). BMC Genomics, 16, 718.

14 Shapulatov,U.M., Buriev,Z.T., Ulloa,M., Saha,S., Devor,E.J., Ayubov,M.S., Norov,T.M., Shermatov,S.E., Abdukarimov,A., Jenkins,J.N., et al. (2016) Characterization of Small RNAs and Their Targets from Fusarium oxysporum Infected and Noninfected Cotton Root Tissues. Plant Mol. Biol. Report., 34, 698–706.

15 Fenton,A., Antonovics,J. and Brockhurst,M.A. (2009) Inverse-Gene-for-Gene Infection Genetics and Coevolutionary Dynamics. Am. Nat., 174, E230–E242.

16 Jones,J.D.G. and Dangl,J.L. (2006) The plant immune system. Nature, 444, 323–329.

17 Roux,F., Voisin,D., Badet,T., Balagué,C., Barlet,X., Huard-Chauveau,C., Roby,D. and Raffaele,S. (2014) Resistance to phytopathogens e tutti quanti: placing plant quantitative disease resistance on the map. Mol. Plant Pathol., 15, 427–432.

18 Asins,M.J. (2002) Present and future of quantitative trait locus analysis in plant breeding. Plant Breed., 121, 281–291.

19 Visscher,P.M., Wray,N.R., Zhang,Q., Sklar,P., McCarthy,M.I., Brown,M.A. and Yang,J. (2017) 10 Years of GWAS Discovery: Biology, Function, and Translation. Am. J. Hum. Genet., 101, 5–22.

20 Brachi,B., Morris,G.P. and Borevitz,J.O. (2011) Genome-wide association studies in plants: the missing heritability is in the field. Genome Biol., 12, 232.

21 Bradbury,P.J., Zhang,Z., Kroon,D.E., Casstevens,T.M., Ramdoss,Y. and Buckler,E.S. (2007) TASSEL: software for association mapping of complex traits in diverse samples. Bioinformatics, 23, 2633–2635.

22 Purcell,S., Neale,B., Todd-Brown,K., Thomas,L., Ferreira,M.A.R., Bender,D., Maller,J., Sklar,P., de Bakker,P.I.W., Daly,M.J., et al. (2007) PLINK: A Tool Set for Whole-Genome Association and Population-Based Linkage Analyses. Am. J. Hum. Genet., 81, 559–575.

23 Zhang,Z., Ersoz,E., Lai,C.-Q., Todhunter,R.J., Tiwari,H.K., Gore,M.A., Bradbury,P.J., Yu,J., Arnett,D.K., Ordovas,J.M., et al. (2010) Mixed linear model approach adapted for genome-wide association studies. Nat. Genet., 42, 355–360.

24 Bergelson,J. and Roux,F. (2010) Towards identifying genes underlying ecologically relevant traits in Arabidopsis thaliana. Nat. Rev. Genet., 11, 867–879.

25 Huard-Chauveau,C., Perchepied,L., Debieu,M., Rivas,S., Kroj,T., Kars,I., Bergelson,J., Roux,F. and Roby,D. (2013) An Atypical Kinase under Balancing Selection Confers Broad-Spectrum Disease Resistance in Arabidopsis. PLoS Genet., 9, e1003766.

26 Badet,T., Voisin,D., Mbengue,M., Barascud,M., Sucher,J., Sadon,P., Balagué,C., Roby,D. and Raffaele,S. (2017) Parallel evolution of the POQR prolyl oligo peptidase gene conferring plant quantitative disease resistance. PLOS Genet., 13, e1007143.

27 Navaud,O., Barbacci,A., Taylor,A., Clarkson,J.P. and Raffaele,S. (2018) Shifts in diversification rates and host jump frequencies shaped the diversity of host range among Sclerotiniaceae fungal plant pathogens. Mol. Ecol., 27, 1309–1323.

28 Kabbage,M., Yarden,O. and Dickman,M.B. (2015) Pathogenic attributes of Sclerotinia sclerotiorum: Switching from a biotrophic to necrotrophic lifestyle. Plant Sci., 233, 53–60.

29 Derbyshire,M.C. and Denton-Giles,M. (2016) The control of sclerotinia stem rot on oilseed rape (Brassica napus): current practices and future opportunities. Plant Pathol., 65, 859–877.

30 Derbyshire,M., Denton-Giles,M., Hegedus,D., Seifbarghy,S., Rollins,J., van Kan,J., Seidl,M.F., Faino,L., Mbengue,M., Navaud,O., et al. (2017) The Complete Genome Sequence of the Phytopathogenic Fungus Sclerotinia sclerotiorum Reveals Insights into the Genome Architecture of Broad Host Range Pathogens. Genome Biol. Evol., 9, 593–618.

31 Badet,T., Peyraud,R., Mbengue,M., Navaud,O., Derbyshire,M., Oliver,R.P., Barbacci,A. and Raffaele,S. (2017) Codon optimization underpins generalist parasitism in fungi. Elife, 6.

32 Bolger,A.M., Lohse,M. and Usadel,B. (2014) Trimmomatic: a flexible trimmer for Illumina sequence data. Bioinformatics, 30, 2114–2120.

33 Formey,D., Iñiguez,L.P., Peláez,P., Li,Y.-F., Sunkar,R., Sánchez,F., Reyes,J.L. and Hernández,G. (2015) Genome-wide identification of the Phaseolus vulgaris sRNAome using small RNA and degradome sequencing. BMC Genomics, 16, 423.

34 Anders,S. and Huber,W. (2010) Differential expression analysis for sequence count data. Genome Biol., 11, R106.

35 Dai,X. and Zhao,P.X. (2011) psRNATarget: a plant small RNA target analysis server. Nucleic Acids Res., 39, W155–W159.

36 Jurka,J., Kapitonov,V.V., Pavlicek,A., Klonowski,P., Kohany,O. and Walichiewicz,J. (2005) Repbase Update, a database of eukaryotic repetitive elements. Cytogenet. Genome Res., 110, 462–467.

37 Alexa,A., Rahnenfuhrer,J. and Lengauer,T. (2006) Improved scoring of functional groups from gene expression data by decorrelating GO graph structure. Bioinformatics, 22, 1600–1607.

38 Benjamini,Y. and Hochberg,Y. (1995) Controlling the False Discovery Rate: A Practical and Powerful Approach to Multiple Testing. J. R. Stat. Soc. Ser. B, 57, 289–300.

39 Coolen,S., Proietti,S., Hickman,R., Davila Olivas,N.H., Huang,P.-P., Van Verk,M.C., Van Pelt,J.A., Wittenberg,A.H.J., De Vos,M., Prins,M., et al. (2016) Transcriptome dynamics of Arabidopsis during sequential biotic and abiotic stresses. Plant J., 86, 249–267.

40 Hooton,J.W.L. (1991) Randomization tests: statistics for experimenters. Comput. Methods Programs Biomed., 35, 43–51.

41 Wang,M., Weiberg,A., Lin,F.-M., Thomma,B.P.H.J., Huang,H.-D. and Jin,H. (2016) Bidirectional cross-kingdom RNAi and fungal uptake of external RNAs confer plant protection. Nat. Plants, 21, 16151.

42 Yang,F. (2015) Genome-wide analysis of small RNAs in the wheat pathogenic fungus Zymoseptoria tritici. Fungal Biol., 119, 631–640.

43 Vetukuri,R.R., Åsman,A.K.M., Tellgren-Roth,C., Jahan,S.N., Reimegård,J., Fogelqvist,J., Savenkov,E., Söderbom,F., Avrova,A.O., Whisson,S.C., et al. (2012) Evidence for Small RNAs Homologous to Effector-Encoding Genes and Transposable Elements in the Oomycete Phytophthora infestans. PLoS One, 7, e51399.

44 Ghildiyal,M., Xu,J., Seitz,H., Weng,Z. and Zamore,P.D. (2010) Sorting of Drosophila small silencing RNAs partitions microRNA* strands into the RNA interference pathway. RNA, 16, 43–56.

45 Mi,S., Cai,T., Hu,Y., Chen,Y., Hodges,E., Ni,F., Wu,L., Li,S., Zhou,H., Long,C., et al. (2008) Sorting of Small RNAs into Arabidopsis Argonaute Complexes Is Directed by the 5′ Terminal Nucleotide. Cell, 133, 116–127.

46 Fang,X. and Qi,Y. (2016) RNAi in Plants: An Argonaute-Centered View. Plant Cell, 28, 272–285.

47 Croll,D. and McDonald,B.A. (2012) The Accessory Genome as a Cradle for Adaptive Evolution in Pathogens. PLoS Pathog., 8, e1002608.

48 Raffaele,S. and Kamoun,S. (2012) Genome evolution in filamentous plant pathogens: why bigger can be better. Nat. Rev. Microbiol., 10, 417–430.

49 Rouxel,T., Grandaubert,J., Hane,J.K., Hoede,C., van de Wouw,A.P., Couloux,A., Dominguez,V., Anthouard,V., Bally,P., Bourras,S., et al. (2011) Effector diversification within compartments of the Leptosphaeria maculans genome affected by Repeat-Induced Point mutations. Nat. Commun., 2, 202.

50 Spanu,P.D., Abbott,J.C., Amselem,J., Burgis,T.A., Soanes,D.M., Stuber,K., Loren van Themaat,E.V., Brown,J.K.M., Butcher,S.A., Gurr,S.J., et al. (2010) Genome Expansion and Gene Loss in Powdery Mildew Fungi Reveal Tradeoffs in Extreme Parasitism. Science (80-.)., 330, 1543–1546.

51 Seidl,M.F. and Thomma,B.P.H.J. (2017) Transposable Elements Direct The Coevolution between Plants and Microbes. Trends Genet., 33, 842–851.

52 Van Kan,J.A.L., Stassen,J.H.M., Mosbach,A., Van Der Lee,T.A.J., Faino,L., Farmer,A.D., Papasotiriou,D.G., Zhou,S., Seidl,M.F., Cottam,E., et al. (2017) A gapless genome sequence of the fungus Botrytis cinerea. Mol. Plant Pathol., 18, 75–89.

53 Syme,R.A., Tan,K.-C., Hane,J.K., Dodhia,K., Stoll,T., Hastie,M., Furuki,E., Ellwood,S.R., Williams,A.H., Tan,Y.-F., et al. (2016) Comprehensive Annotation of the Parastagonospora nodorum Reference Genome Using Next-Generation Genomics, Transcriptomics and Proteogenomics. PLoS One, 11, e0147221.

54 Laurent,B., Moinard,M., Spataro,C., Ponts,N., Barreau,C. and Foulongne-Oriol,M. (2017) Landscape of genomic diversity and host adaptation in Fusarium graminearum. BMC Genomics, 18, 203.

55 Dallery,J.-F., Lapalu,N., Zampounis,A., Pigné,S., Luyten,I., Amselem,J., Wittenberg,A.H.J., Zhou,S., de Queiroz,M.V., Robin,G.P., et al. (2017) Gapless genome assembly of Colletotrichum higginsianum reveals chromosome structure and association of transposable elements with secondary metabolite gene clusters. BMC Genomics, 18, 667.

56 Guyon,K., Balagué,C., Roby,D. and Raffaele,S. (2014) Secretome analysis reveals effector candidates associated with broad host range necrotrophy in the fungal plant pathogen Sclerotinia sclerotiorum. BMC Genomics, 15, 336.

57 Chen,C.-W., Panzeri,D., Yeh,Y.-H., Kadota,Y., Huang,P.-Y., Tao,C.-N., Roux,M., Chien,S.-C., Chin,T.-C., Chu,P.-W., et al. (2014) RETRACTED: The Arabidopsis Malectin-Like Leucine-Rich Repeat Receptor-Like Kinase IOS1 Associates with the Pattern Recognition Receptors FLS2 and EFR and Is Critical for Priming of Pattern-Triggered Immunity. Plant Cell, 26, 3201–3219.

58 Wen,Z., Yao,L., Wan,R., Li,Z., Liu,C. and Wang,X. (2015) Ectopic Expression in Arabidopsis thaliana of an NB-ARC Encoding Putative Disease Resistance Gene from Wild Chinese Vitis pseudoreticulata Enhances Resistance to Phytopathogenic Fungi and Bacteria. Front. Plant Sci., 6, 1087.

